# Structure and thermodynamics of a UGG motif interacting with Ba^2+^ and other metal ions: accommodating changes in the RNA structure and the presence of a G(*syn*)–G(*syn*) pair

**DOI:** 10.1101/2022.08.11.503657

**Authors:** Agnieszka Kiliszek, Martyna Pluta, Magdalena Bejger, Wojciech Rypniewski

**Affiliations:** Institute of Bioorganic Chemistry, Polish Academy of Sciences, Noskowskiego 12/14, 61-704 Poznan, Poland

**Keywords:** UGG repeats, U·G wobble, G(syn)–G(syn) pair, RNA crystallography, RNA UV melting, RNA DSC measurment

## Abstract

The self-complementary triplet 5’UGG3’/5’UGG3’ is a particular structural motif containing noncanonical G-G pair and two U·G wobble pairs. It constitutes a specific structural and electrostatic environment attracting metal ions, particularly Ba^2+^ ions. Crystallographic research has shown that two Ba^2+^ cations are located in the major groove of the helix and interact directly with the UGG triplet. A comparison with the unliganded structure has revealed global changes in the RNA structure in the presence of metal ions, whereas thermodynamic measurements have shown increased stability. Moreover, in the structure with Ba^2+^, an unusual noncanonical G(*syn*)-G(*syn*) pair is observed instead of the common G(*anti*)-G(*syn*). We further elucidate the metal binding properties of the UGG/UGG triplet by performing crystallographic and thermodynamic studies using DSC and UV melting with other metal ions. The results explain the preferences of the UGG sequence for Ba^2+^ cations and point to possible applications of this metal-binding propensity.

## INTRODUCTION

RNA contains many noncanonical base pairs that increase the structural richness of RNA compared to fully complementary DNA. Due to their unsaturated potential for base pairing, these noncanonical pairs have an additional capacity for binding external ligands, including metal cations. These bound cations play a role in the folding of RNA into tertiary structures and can also be involved in catalysis conducted by rybozymes.

The G·U wobble is the most common noncanonical base pair found in RNA structures (Varani and Mcclain 2000). In most cases the G·U pair is stabilized by two hydrogen bonds: N1···O2 and O6···N3. The guanosine residue is inclined toward the minor groove, while the uridine residue is shifted toward the major groove. This arrangement results in the exposition of functional groups (N2 *exo*-amino of G, O4 carbonyl of U) and the formation of a specific topological and electrostatic landscape within the RNA helix. The functional groups not involved in G·U pairing show a high binding potential for ligands (Strobel and Cech 1995, 1996; Ananth et al. 2013). The O4 carbonyl group of uridine together with the N7 and O6 atoms of guanine creates a broad electronegative potential (Xu et al. 2007). This attracts metal ions and other positively charged ligands (Allain and Varani 1995; Burgstaller 1997; Ananth et al. 2013).

X-ray and NMR studies have provided much information about structures containing either single or tandem G·U wobbles and enriched our knowledge about the properties of this noncanonical base pair (Mooers et al. 1997; Mueller et al. 1999; Trikha et al. 1999; Sashital et al. 2007; Ruszkowska et al. 2022; McDowell et al. 1997; Gu et al. 2015; Garg and Heinemann 2018; Biswas and Sundaralingam 1997; Berger et al. 2019). An important feature of the G·U pair is that it is nonisosteric, i.e., the superposition of N-glycosic bonds of 5’G·U3’ with the 5’U·G3’ pair results in a large shift between the bases (Westhof et al. 2019). The most pronounced outcome of the nonisostericity is observed in the twist angle between G·U and the neighboring pair. For G·U, the following pair is usually undertwisted while for U·G it is overtwisted (Ananth et al. 2013). Overall, the twist is distinct from a typical value observed for A-RNA but the overall effect on the helix structure is moderate and absorbed by local changes in the sugar–phosphate backbone.

There are several known three-dimensional models of a wobble pair with bound metal ions in a major groove (Ruszkowska et al. 2022; Price et al. 2015; Keel et al. 2007; Kacer et al. 2003; Colmenarejo and Tinoco 1999; Allain and Varani 1995; Gu et al. 2015). Structural studies have confirmed the ability of G·U to interact with metal species but have also indicated that the potential of the wobble motif to bind cations depends on their structural context (Keel et al. 2007). The structural and electrostatic requirements for the metal ions to bind this wobble pair have not yet been fully investigated. Here, we present crystallographic and thermodynamic studies of the self-complementary triplet 5’UGG3’/5’UGG3’ motif consisting of two U·G wobble pairs interrupted by a G-G pair. The presented motif can be found in humans within the mRNA of certain genes as a repeated sequence. Intronic UGG repeats participate in alternative splicing of the *CD44* gene that codes the CD44 antigen involved in cell–cell interactions, adhesion and migration (Galiana-Arnoux et al. 2005). The crystallographic structures reveal that the UGG motif shows particular affinity for Ba^2+^ forming direct interactions with bound metal ions. The presence of Ba^2+^ is associated with global changes in RNA structure and with the presence of an unusual noncanonical G(*syn*)-G(*syn*) pair instead of the common G(*anti*)-G(*syn*). To elucidated the metal binding properties of the UGG/ UGG triplet, we perform crystallographic and thermodynamic studies using DSC and UV melting with other metal ions. The results explain the preferences of the UGG sequence for Ba^2+^ cations. These properties can be useful in crystallographic research, for phasing with anomalous signal, and in biotechnology, whether using RNA molecules as a metal sponge or as a sensor of metal ions.

## RESULTS

### The RNA models

Seven crystal structures of the RNA duplex (AUG<u>UGG</u>CAU)_2_ containing the 5’UGG3’/5’UGG3’ motif were determined: in an unliganded form and in the presence of the following metal cations: Cs^+^, Sr^2+^, Ca^2+^, Cd^2+^ and two models with Ba^2+^ ions (Table S1). All six of the ion-liganded structures were crystallized in similar conditions, containing MPD as the precipitant, while the unliganded structure crystallized in the presence of ammonium sulfate. The Ba^2+^ ions were identified as strong peaks (approx. 14 r.m.s.d.) on the anomalous electron density map, and confirmed by an examination of their coordination spheres and distances from ligand molecules. Crystals containing Cs^+^, Sr^2+^, Ca^2+^ and Cd^2+^ were obtained by the cocrystalization of RNA with the appropriate ions, in the absence of Ba^2+^. In all four structures the RNA molecules showed the same packing in the crystal lattice resulting in the same space group and unit cell parameters. The asymmetric unit of the monoclinic crystals contained fourteen independent RNA duplexes. A weak anomalous signal was also observed for Cs^+^ and Sr^2+^ structures.

### Noncanonical pairs in RNA duplexes

The AUG<u>UGG</u>CAU oligomer forms self-complementary duplexes. The six flanking nucleotides are engaged in Watson–Crick base pairing between complementary nucleotides of the two strands (Figure 1A), with the exception of the RNA-Ba(1) structure, in which the 1A residue of chain C and the 9U residue of chain D form a noncanonical pair (see details in Supplemental Material Figure S1). The UGG motif is located in the middle of the duplex and is engaged in noncanonical pairing. Two U·G pairs are wobbled, and the G–G pairs are in most cases G(*syn*)– G(*anti*), but in the RNA-Ba(2) structure there is also an instance of the G(*syn*)–G(*syn*) conformation (Figure 2A-D). The G(*syn*)–G(*anti*) pairs interact *via* two H-bonds (N1···O6 and N2···N7) between their respective Watson–Crick and Hoogsteen edges, as described earlier (Figure 2B) (Kiliszek et al. 2011). The G(*syn*)–G(*syn*) pair has not been described previously. The G(*syn*) bases interact *via* two solvent molecules wedged between them. The innermost ligand is 2.2–2.5 Å from the carbonyl O6 atoms and the other ligand. Its distance from the N7 atoms is 2.5–2.8 Å. The other ligand is 2.4–2.6 Å from the two N7 atoms. The closeness of the inner ligand to the carbonyl O2 atoms suggests that it could be Na^+^ rather than a water molecule, while the outer ligand could be water-donating H-bonds to the imino groups (Figure 2D). The unusual alignment of the G rings is stabilized by the presence of two Ba^2+^ ions in the major groove at 3.0–3.1 Å from the carbonyl groups (Figure 1B and 3A). Indications of G(*syn*)–G(*syn*) pairs are also observed in RNA-Cs, RNA-Cd and RNA-Ba(1) structures but the modeling is ambiguous due to static disorder and low resolution.

**Figure 1.**
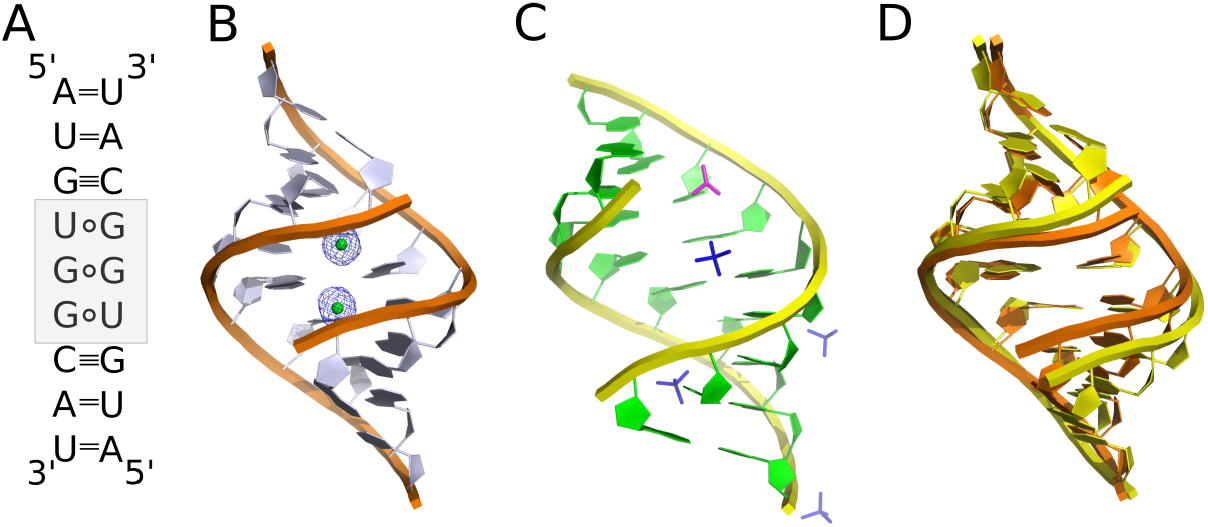
Secondary and tertiary structure models of the AUG<u>UGG</u>CAU duplex. (A) The UGG/UGG motif (*gray*) forms two U·G pairs and one noncanonical G-G pair. (B) Cartoon representation of the RNA-Ba(2) model with bound Ba^2+^ ions (*green balls*) in the major groove of the helix. The 2F_o_–F_c_ electron density map (*gray*) is contoured at the 1σ level. (3) Cartoon representation of the unliganded model with sulfate (*dark blue sticks*) and acetate ions (*pink sticks*) located in the major and minor grooves of the helix. (D) A superposition of RNA-Ba(2) (*orange*) and unliganded structures (*yellow*).

**Figure 2.**
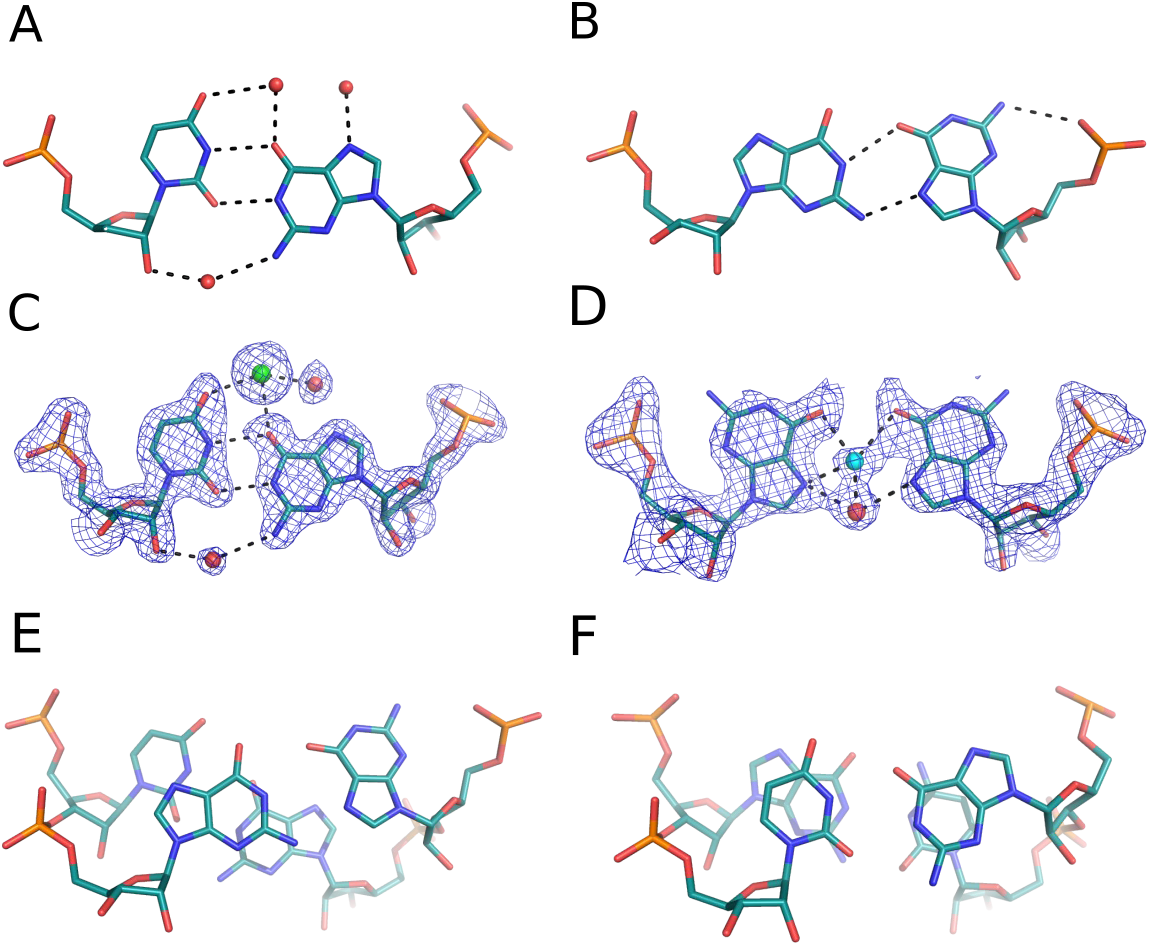
Noncanonical base pairs observed in crystallographic models of the AUG<u>UGG</u>CAU duplex. (A and C) The U·G wobble interacting with solvent molecules. (B) Stick representation of the G(*anti*)–G(*syn*) pair and (D) G(*syn*)–G(*syn*) base pairs. The stacking interactions observed for G-G (E) and U·G pairs (F). Water molecules are red spheres, Ba^2+^ ions are green, and Na^+^ is blue. The 2F_o_–F_c_ electron density map (gray) is contoured at the 1σ level. The H-bonds are black dashed lines.

**Figure 3.**
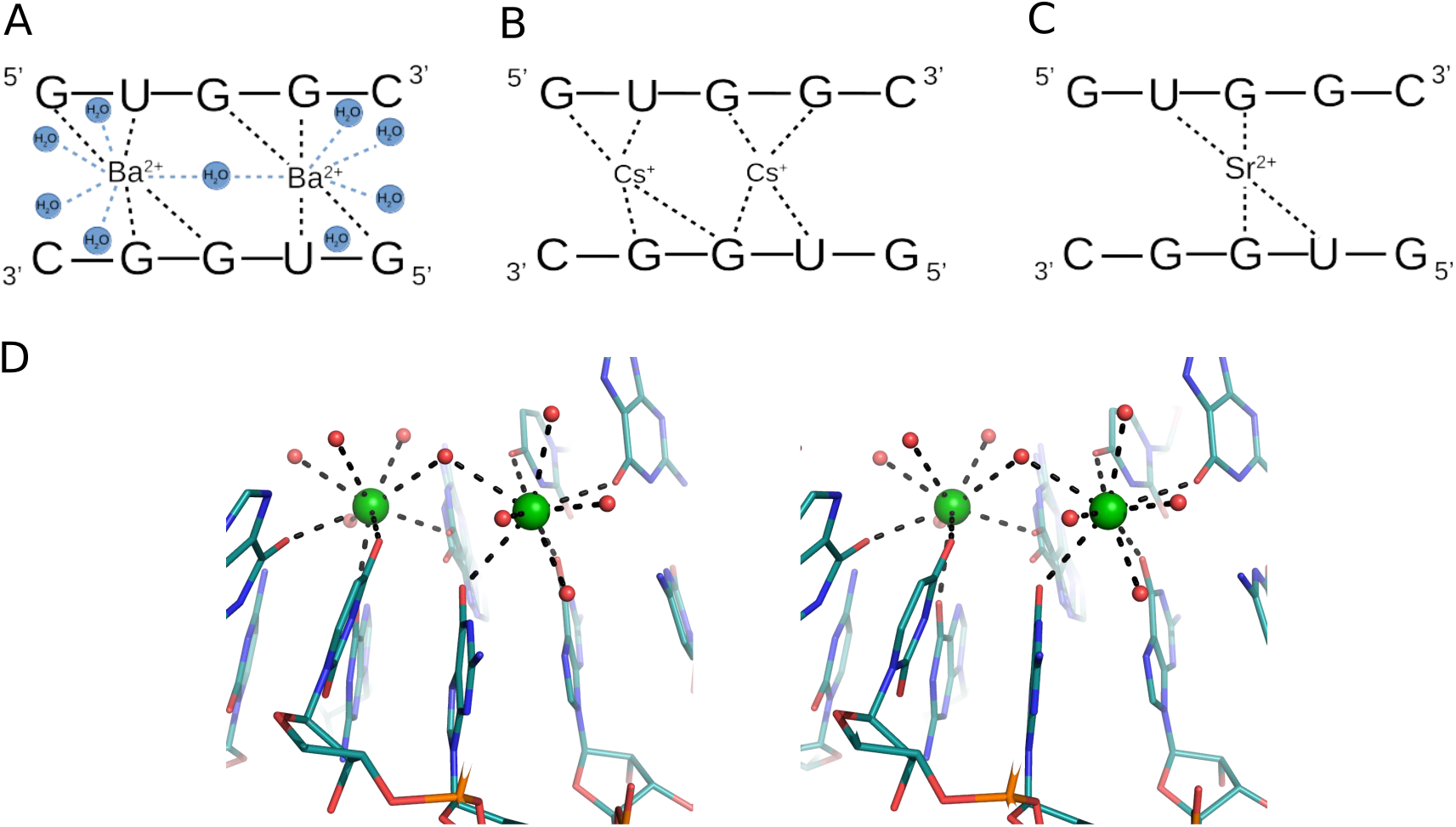
Interactions of RNA residues with metal ions: (A) with Ba^2+^, (B) Cs^+^ and (C) Sr^2+^ ions. (D) Stereo view of RNA interacting with Ba^2+^ ions (*green spheres*).

### Interactions of RNA with metal ions

Each UGG motif can bind up to two metal ions in the major groove. Complexes with Ba^2+^, Cs^+^ and Sr^2+^ are formed easily, while Ca^2+^ and Cd^2+^ are difficult to observe. Ba^2+^ and Cs^+^ occupy two binding sites, while Sr^2+^ shows a mixture of binding modes (Figure 1B, and 3A–C; Figure S2). In seven out of fourteen crystallographically independent duplexes single Sr^2+^ ions are bound; in three cases there are two ions while in four cases, the occupancy of the second ion is only partial (Figure S2). Single Cd^2+^ ions are observed in only three duplexes. In the structure crystallized with Ca^2+^, the solvent structure is indistinct. Three single peaks of the electron density map observed in the major groove have been interpreted as water molecules, not Ca^2+^ (Figure S2). Their coordination distances are at least 2.7 Å, while for calcium, they should be approx. 2.4 Å (Harding et al. 2010).

When two ions (Ba^2+^, Cs^+^ or Sr^2+^) interact with the UGG motif, they are located on each side of the G-G pair, above the U·G wobble (Figure 3A and 3B). For Ba^2+^ the pattern of interactions is the most ordered. Each ion forms four direct interactions with the carbonyl groups of RNA, spanning both strands of the duplex. Two bonds are observed with O6 of 3G and O4 of 4U residues of one chain and with O6 of 5G and O6 of 6G of the second strand. In addition, Ba^2+^ interacts with\ up to five water molecules, supplementing its coordination sphere to a maximum value of nine ligands. The range of distances is 2.5–3.1 Å. The ions are approx. 4.8 Å apart and share only one water molecule (Figure 3A and 3D). In the case of Cs^+^, the pattern of interactions is less ordered. Each Cs^+^ ion forms four direct contacts with RNA, as in the case of Ba^2+^, but in some cases, one of the ions is shifted toward the G-G pair engaging the carbonyl groups of both guanosines (Figure 3B). This is reflected in a wide range of Cs-Cs distances, from 4.8 to 5.7 Å. Similarly, in the Sr-structure, one of the metal ions tends to be shifted toward the G-G pair. When only one ion (Sr^2+^ or Cd^2+^) is bound to RNA, it is placed just above the G-G pair interacting with their carbonyl groups. Two additional contacts can be observed with the carbonyl groups of the 4U residues of each strand (Figure 3C).

In the metal-free UGG structure, water molecules are observed in the major groove of the U·G wobble. Each water molecule interacts with U·G and the neighboring G(*syn*)–G(*anti*) pair (Figure 2A). In addition to water, four sulfate and one acetate ions have been identified, either in the major or the minor groove, interacting mostly with positively charged amino groups of RNA (Figure 1C).

### Impact of metal ions on the RNA structure

In all the described models the RNA has the A-form (Figure 1B–C and Figure S2). The presence of the noncanonical base pairs seems to have a local rather than global effect on the structure. The conformation of the U•G and G-G pairs is reflected in the shear and shift parameters. Both have distinct values from the neighboring pairs (Table S2 and S3). Additionally, the arrangement of the two bulky G-G residues is reflected in the stretch parameter. Other parameters, such as helical twist, groove width or propeller are associated with the presence of metal ions. A comparison of the unliganded structure with Ba-models revealed that in the presence of Ba^2+^, the major groove is narrowed, and access to it is limited (Figure 1D). The major groove width for the native model is 14.2 Å, while for Ba-RNA the values are in the range of 9.7–10.8 Å. In the other complexes with metal ions, groove narrowing is not observed. Rather, they exhibit a range of groove widths (10.1–15.7 Å). The twist parameter is correlated with the groove width. For the Ba-models, the twist has the highest value (34.5°±5.2°), while for other metal-structures, it is 33.3°±3.0°, and for the unliganded form, it is 32.1°±3.7° (Table S4). The other parameter associated with Ba^2+^ ions is the propeller twist which seems to be elevated (16.9°–23.0°) in comparison to the native structure (14.0°) and other metal-structures (less than 17°) (Table S5).

### Stacking interactions and electrostatic profile distribution

The distribution of stacking interactions in all the models is similar. Since each duplex possesses self-symmetry, the stacking is symmetric. Intrastrand overlaps are predominant. Only for the 2U3G/7C8A steps, is an interstack observed between the guanosine and adenosine residues. Most extensive stacking interactions are observed between the neighboring 3G4U and 6G7C residues. Measuring only the overlap of base rings (excluding exocyclic atoms) the area is on average 4 Å^2^, which covers almost all of the pyrimidine plane (Figure 2F). The observed distribution of stacking indicates that the 4U and 6G residues involved in wobble pairs interact extensively, while guanosines forming the G–G pairs are engaged least of all the nucleotides in the duplex (Figure 2E-F).

All the helices exhibit similar electrostatic potential and surface profile. The major groove is mostly electronegative due to the presence of carbonyl groups. Patches of positive potential come from the exposed Watson–Crick 5G(*syn*) edge and the *exo*-amino group of 8C residues (Figure 4A and 4C). The G–G pairs are located in the center of the helix and form a platform of positive and negative potential while the neighboring wobble pairs are strictly electronegative. The guanosine from the U•G pair is pushed toward the minor groove and forms a niche, but it does not seem to be occupied, and the metal ions occupy the central part of the electronegative area of the wobble pairs (Figure 4C). When only one ion is bound to the RNA, it is located in the center of the helix above the negatively charged region of the G-G pair. The minor groove shows a characteristic RNA electrostatic profile with alternating stripes of positive and negative potential. The *exo*-amino group of the wobbled guanosines seems to be less exposed in comparison to other residues (Figure 4B and 4D).

**Figure 4.**
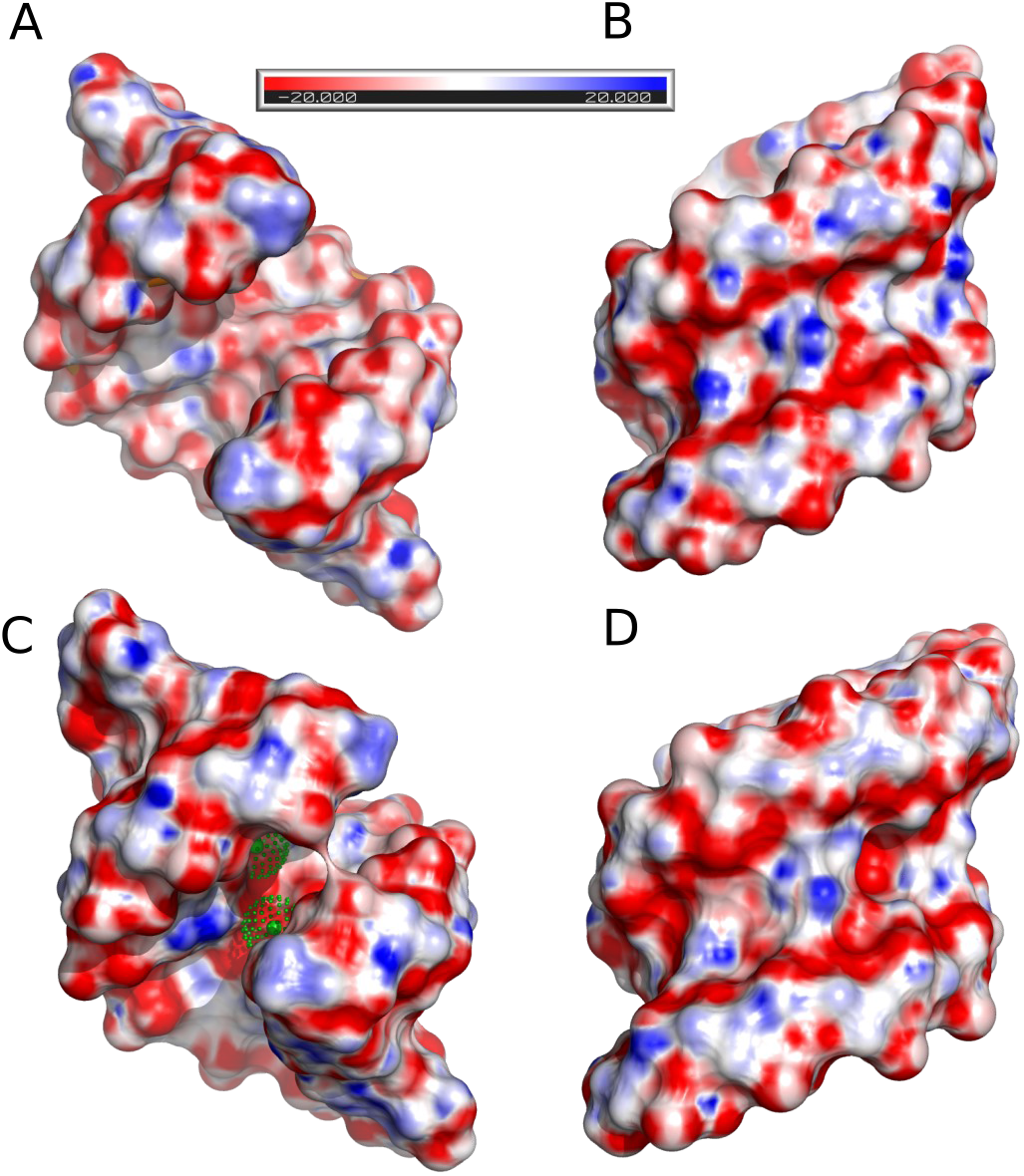
The electrostatic potential distribution calculated for (A and B) the unliganded structure and (C and D) the RNA-Ba(2) model. The Ba^2+^ ions are green spheres.

### Thermal stability of RNA

Thermodynamic and thermal studies were performed for the AUG<u>UGG</u>CAU duplex in the presence of Cs^+^, Sr^2+^, Ca^2+^, Cd^2+^ or Ba^2+^ ions using the UV melting method and differential scanning calorimetry (DSC). The measurements were performed in 100 mM NaCl and 5 mM of metal ions, since the same conditions were present in the crystallization solutions. The reference RNA sample was melted in the buffer containing only 100 mM NaCl. Additionally, the thermal stability of the duplex was determined in high salt concentrations: (1) 1 M NaCl and (2) 1 M NaCl with 5 mM BaCl_2_. The UV-melting data were used to calculate thermodynamic parameters such as ΔH, ΔG and ΔS and to determine the melting temperature T_M_ of the duplex at 100 µM RNA concentration (Table 1 and Figure S3). Each DSC measurement was performed in five runs of the denaturation– renaturation cycle to define the melting point of the duplex T_m_ (as distinct from T_M_ which was calculated for the UV-melting experiments), the degree of degradation or refolding of RNA and the presence of other forms of RNA aggregation (Table 1 and Table S6).

**Table 1.** The thermodynamic and thermal papameters of the AUGUGGCAU oligomer in different buffer condition.

| Buffer conditions | Analysis of Melt Curve Fits/Errors | | | | Analysis of $T_m$ Dependence/Errors | | | | DSC | | |
| --- | --- | --- | --- | --- | --- | --- | --- | --- | --- | --- | --- |
| | $-\Delta H^\circ$ (kcal/mol) | $-\Delta S^\circ$ (eu) | $-\Delta G^\circ_{37}$ (kcal/mol) | $T_m$ (°C) | $-\Delta H^\circ$ (kcal/mol) | $-\Delta S^\circ$ (eu) | $-\Delta G^\circ_{37}$ (kcal/mol) | $\Delta\Delta G^\circ_{37}$ (kcal/mol) | $T_m$ (°C) | $T_m$ (°C) at given RNA concentration | Degree of RNA degradation |
| 0.1 M NaCl | $49.8 \pm 4.8$ | $151.1 \pm 16.3$ | $2.90 \pm 0.30$ | 20.8 | $55.0 \pm 5.3$ | $69.0 \pm 18.1$ | $2.60 \pm 0.37$ | --- | 20.6 | 24.6 (300μM) | low |
| BaCl<sub>2</sub>,<br>0.1 M NaCl | $63.2 \pm 2.7$ | $184.6 \pm 8.6$ | $5.94 \pm 0.12$ | 38.3 | $66.5 \pm 3.3$ | $195.4 \pm 10.8$ | $5.91 \pm 0.05$ | 3.31 | 38.1 | 40.6 (260μM) | medium |
| SrCl<sub>2</sub>,<br>0.1 M NaCl | $62.3 \pm 3.0$ | $187.6 \pm 9.6$ | $4.09 \pm 0.33$ | 29.3 | $66.9 \pm 13.4$ | $203.1 \pm 44.4$ | $3.92 \pm 0.67$ | 1.32 | 29.1 | 34.1 (291μM) | medium |
| CaCl<sub>2</sub>,<br>0.1 M NaCl | $55.1 \pm 10.0$ | $165.7 \pm 33.7$ | $3.73 \pm 0.54$ | 26.4 | $46.2 \pm 4.9$ | $135.0 \pm 16.5$ | $4.28 \pm 0.27$ | 1.68 | 27.9 | 32.7 (211μM) | high |
| CsCl,<br>0.1 M NaCl | $58.6 \pm 3.0$ | $180.9 \pm 10.1$ | $2.51 \pm 0.15$ | 21.1 | $62.6 \pm 6.6$ | $194.6 \pm 22.6$ | $2.24 \pm 0.45$ | -0.36 | 20.8 | 23.6 (251μM) | no degradation |
| CdCl<sub>2</sub>,<br>0.1 M NaCl | $32.4 \pm 2.1$ | $92.3 \pm 6.5$ | $3.81 \pm 0.11$ | 20.1 | $46.4 \pm 8.1$ | $140.6 \pm 28.2$ | $2.75 \pm 0.65$ | 0.15 | 18.6 | 21.8 (160μM) | high |
| 1 M NaCl | $56.1 \pm 1.5$ | $165.5 \pm 5.0$ | $4.77 \pm 0.07$ | 32.1 | $60.3 \pm 1.1$ | $179.5 \pm 3.6$ | $4.64 \pm 0.03$ | 2.04 | 31.8 | 36.0 (221μM) | low |
| BaCl<sub>2</sub>,<br>1 M NaCl | $50.0 \pm 1.7$ | $144.7 \pm 5.2$ | $5.15 \pm 0.25$ | 33.8 | $74.8 \pm 7.0$ | $226.6 \pm 23.1$ | $4.55 \pm 0.21$ | 1.95 | 32.4 | -- | --- |

Both methods show similar effects on the thermal stability of the investigated samples. The RNA is stabilized the most in the presence of Ba^2+^ (Table 1). The ΔG is 3.3 kcal/mol lower thant that of the reference sample. This amounts to a higher melting temperature by 17.6 °C for UV melting and by 16 °C for DSC measurement. Sr^2+^ and Ca^2+^ seem to have a similar stabilizing effect. For Sr^2+^, it is 1.3 kcal/mol and 8.5 °C or 9.5 °C (UV melting and DSC, respectively), and in the presence of Ca^2+^, it is 1.7 kcal/mol and 7.3 °C or 8.1 °C (UV melting, DSC). Cs^+^ and Cd^2+^ do not stabilize the duplex or show only a slight destabilizing effect. In the presence of Cd^2+^, the RNA has a lower melting temperature by approx. 2 °C and ΔΔG = −0.2 kcal/mol. In the presence of Cs^+^, the melting temperature is similar to that of the control sample, while ΔG is higher by 0.4 kcal/mol. In 1 M NaCl the T_m_ of the duplex is higher than that in 0.1 M NaCl by 11 °C and ΔG is lower by 2 kcal/mol. These parameters indicate that a high concentration of NaCl stabilizes the RNA substantially but not as much as Ba^2+^ ions. When the duplex is melted in 1M NaCl and 5 mM Ba^2+^ the stabilization effect of the metal ions is not observed (the same ΔG for both conditions: 1 M NaCl and 5 mM Ba^2+^ in 1 M NaCl).

In all cases, the DSC spectra show an additional broad peak at higher temperatures (45–90 °C) (Figure 5). It is present only in the first denaturation–renaturation cycle. The height and width of the peak were different for each RNA preparation. In Ba^2+^, the high-temperature peak is the smallest compared to the first peak, while for Cd^2+^, it is relatively the highest peak. In the presence of Sr^2+^, a third peak can be distinguished at 49 °C. The DSC experiments show RNA degradation with each cycle, indicated by diminishing heights of the main peak. The fastest loss of the signal is observed for Ca^2+^ and Cd^2+^ ions, while the highest protection of RNA is observed for Cs^+^. In NaCl, RNA degrades relatively slowly while in the presence of Ba^2+^ and Sr^2+^, the RNA is hydrolyzed at an intermediate rate (Figure 5).

**Figure 5.**
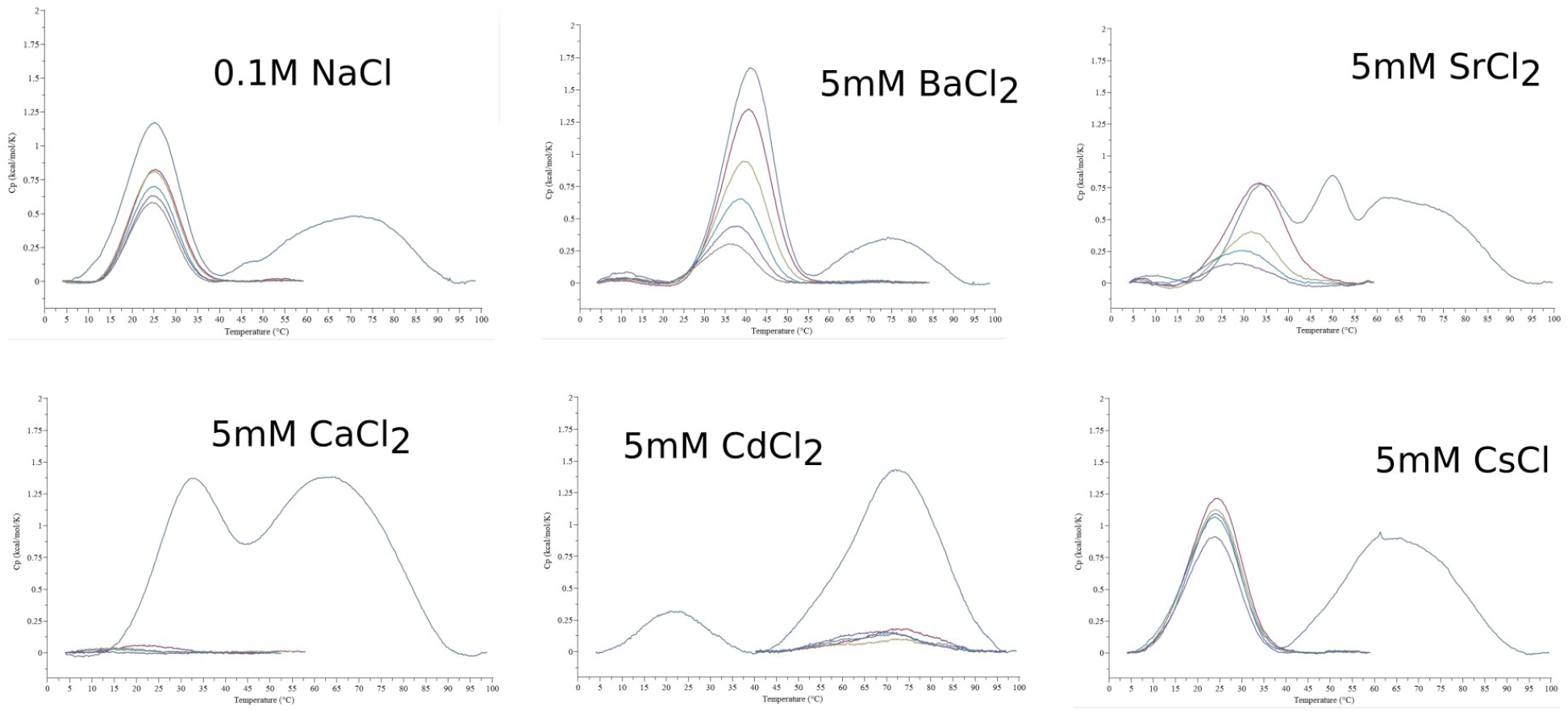
DSC spectra obtained for the AUG<u>UGG</u>CAU oligomer in different buffer conditions.

## DISCUSSION

UGG is interesting from the perspective of structural biology as a motif containing noncanonical pairs G·U and G–G or as a component of microsatellite repeats, but it also plays a biological role. UGG repeats are found in the transcripts of certain genes, mostly in the 3’UTRs (Kozlowski et al. 2010). Intronic UGG repeats participate in alternative splicing of the *CD44* gene which encodes the CD44 antigen involved in cell–cell interactions, adhesion and migration (Galiana-Arnoux et al. 2005).

UGG repeats are noted for forming tetraplexes (Sobczak et al. 2010; Malgowska et al. 2014). A question then arises: why do all the structures in this study have the duplex form? It seems that the crucial variables are the nature of the cations and ionic strength. Tetraplexes have been observed to form mainly in the presence of K^+^, whereas the duplexes presented here contain divalent cations (Ba^2+^, Sr^2+^, Ca^2+^, Cd^2+^) or the monovalent Cs^+^, which stands out, however, as the largest ionic species under study. In the absence of the metal cations, the duplex could be crystallized under the conditions of high ionic strength (2.5 M ammonium sulfate). Another factor favoring the duplex could be the presence of flanking sequences.

Several RNA crystal structures show Ba^2+^ binding at sites consisting of a single UG pair and carbonyl oxygen atoms from neighboring bases, preferably on the 5’-sides (Supplementary Table S7) (Hancock et al. 2004; Daldrop and Lilley 2013; Huang et al. 2016; Price et al. 2015; Schroeder et al. 2012; Huang et al. 2010; Serganov et al. 2009; Bachas and Ferré-D’Amaré 2018; Knappenberger et al. 2018; Heber et al. 2019; Kacer et al. 2003; Ruszkowska et al. 2022). The structure presented in this paper can bind up to two metal ions. The site consists of the 5’UGG/3’GGU motif and two carbonyl oxygen atoms from guanosine rings at the 5’-side of each strand (Figure 3A and 3D). The two cation binding sites are asymmetric due to the asymmetry of the central G(*syn*)–G(*anti*) pair, and they seem to have different affinities for the ions. The exception is the previously unobserved instance of G(*syn*)–G(*syn*), which is symmetric and binds the ligands symmetrically. In the strong complexes, both cation sites are occupied, whereas in the weaker complexes, there can be only one metal ion per UGG motif. The complexes with Ba^2+^ stand out as the best ordered, showing detailed coordination spheres of the metal ions (Figure 3A). Each ion has four RNA ligands and up to five coordinating water molecules. The observed coordination number of 9 is the most common number observed for barium complexes in the CCDC structural database (Hancock et al. 2004). In the complex with Cs^+^, there are also two cations in each UUG. The structure with Sr^2+^ shows a tendency to bind only one metal ion per UGG, and the site is shifted toward the central G–G pair. Cd^2+^ is barely seen, with single sites located in three out of 14 duplexes, while in the structure soaked in Ca^2+^, the presence of the cations is unclear. The apparent tendency to weaker binding and shift toward the G–G pair can be correlated with the decreasing radii of the doubly charged metal ions (Ba^2+^ > Sr^2+^ > Ca^2+^ > Cd^2+^) (Shannon, 1976). This series is reproduced in calorimetric measurements in terms of decreasing thermal stability of the complexes.

Temperature-melting curves and the thermodynamic parameters derived from them were used to determine the amount of stabilization bestowed by the ions on the RNA in solution (Table 1). Ba^2+^ stabilized the most, followed by Sr^2+^ and Ca^2+^, while Cs^+^ and Cd^2+^ had no stabilization effect. The thermodynamic results corroborate the crystallographic observations in terms of their ability to form well-defined complexes, with the exception of Cs^+^ ions, which are clearly visible in the crystal structure but have no impact on the thermal stability of the RNA duplex. They do, however, protect the RNA from hydrolysis compared to all the other examined cations.

How can Cs^+^ shield the chemical integrity of RNA without enhancing its thermal stability? The lack of stabilizing effect could mean that the addition of 5 mM Cs^+^ does not make a significant contribution to stability in comparison to the already present 100 mM Na^+^. Both cations belong to the same group in the periodic table of elements and could play similar roles, balancing the negative charge of the RNA. However, Cs^+^ is much larger, which makes it easier to observe in the crystal structure, and it could fit the major groove more snugly. On the other hand, the protection by Cs^+^ against RNA hydrolysis could mean that this large, singly charged cation does not polarize water efficiently for nucleophilic attack.

Another observation from the thermodynamic study is that the stabilizing effect of Ba^2+^ (ΔΔG=3.0 kcal/mol) is seen only at low ionic strength (0.1 M NaCl), whereas in the presence of 1 M NaCl the amount of stabilization due to Ba^2+^ is much lower (ΔΔG=0.4 kcal/mol) (Table 1). This can be compared with the results of crystallization experiments in which all the complexes with metal ions were crystallized only under low salt conditions (80 mM NaCl), whereas the unliganded RNA could only be crystallized in the presence of high salt (2.5 M ammonium sulfate). The bulk effect of high salt can stabilize the duplex to a comparable degree to the more specifically acting divalent cations. This apparent effect of ionic strength could be relevant to studies in which high salt conditions are used as a point of reference.

Among the RNA motifs containing UG pairs, the structure presented here seems to be an enhanced version of related motifs in terms of its ability to bind metal cations, especially Ba^2+^. It seems that the central G–G pair increases the number of bound Ba^2+^ to two. The affinity for Ba^2+^ is indicated by the well–ordered crystal structures and the amount of thermodynamic stabilization. A distinguishing feature of the RNA complex with Ba^2+^, compared to other cations, is the closing of the major groove around Ba^2+^. This could reflect a good site complementarity of Ba^2+^ in terms of its size and coordination number.

Possible points of interest of this RNA sequence could be an application as an ionic “sponge” or even a detector because the closing of the major groove could be used as a means to activate suitable fluorophores attached to the opposite ends of the motif. A more obvious application could be the use of bound Ba^2+^ in crystallographic phasing. Lastly, a cautionary note to the story is that the ability of Ba^2+^ to bind, with high affinity, to some biological molecules could be a risk factor in the use of barium salts in medical imaging.

## MATERIAL AND METHODS

### Synthesis, purification and crystallization of the AUG<u>UGG</u>CAU oligomer

The oligomer was synthesized by the solid phase method in an Applied Biosystems DNA/RNA synthesizer using TOM protected phosphoramidites. The synthesis was carried out in DMT-ON mode resulting in the oligomer carrying a trityl group at the 5’ end. The RNA was purified according to the protocol suitable for Glen-Pak cartridges (Glen Research) and then desalted and lyophilized under vacuum using a Speed-Vac. The purity of the oligomer was monitored using analytical HPLC separation (Lang and Micura 2008). Usually, the purity was at least 90%, which is suitable for crystallization. Before crystallization the RNA was dissolved in 10 mM sodium cacodylate pH 7.0 to a final concentration of 1 mM. The oligomer was denatured for 5 min at 95 °C and cooled to ambient temperature within 2 hours. Crystals were grown by the sitting drop method at 19 °C. Most of the crystals grew under similar conditions: 0.08 M NaCl, 0.04 M sodium cacodylate pH 6.0, 45% MPD and 12 mM spermine. Depending on the metal ions the solution was supplemented with 20 mM BaCl_2_ or CdCl_2_ or SrCl_2_ or CaCl_2_ or CsCl. Only the unliganded structure was crystallized in 10 mM magnesium acetate, 50 mM MES pH 5.6 and 2.5 M ammonium sulfate. The RNA was mixed with the crystallization solution at a ratio of 3:1.

### X-ray data collection, structure solution, and refinement

X-ray diffraction data were collected on the BL 14.1, 14.2 and 14.3 beam lines at the BESSY II electron storage ring operated by Helmholtz-Zentrum Berlin (Mueller et al. 2015). Only the unliganded crystals were cryoprotected by 20% glycerol (v/v) in the mother liquor. The data were integrated and scaled using XDSAPP or XDSGUI software implementing the XDS package (Kabsch 2010; Krug et al. 2012). The X-ray data are summarized in Table S1. First, the structure of RNA-Ba(1) was solved by anomalous phasing. Although the data were collected at λ=0.895 wavelength Å, far from the absorption edge of barium, the anomalous signal was observed up to a resolution of 3.4 Å. The initial phases were calculated using the Auto-Rickshaw webserver (Panjikar et al. 2009). An inspection of the electron density map indicated that it corresponded well with the determined location of heavy atoms. Moreover, the location of the RNA chain could be resolved. The atomic model was manually built using Coot (Emsley and Cowtan 2004). The remaining structures were solved by molecular replacement with PHASER using RNA-Ba(1) as a search model (Read et al. 2013). Early stages of the refinement were carried out using Refmac5 from the CCP4 program suite (Winn et al. 2011; Vagin et al. 2004), and then refinement was continued with PHENIX (Liebschner et al. 2019). The models are summarized in Table S1. The helical parameters were calculated with 3DNA (Zheng et al. 2009)using a sequence-independent method based on vectors connecting the C1′ atoms of the paired residues to avoid computational artifacts arising from noncanonical base-pairing. The rogram PBEQ-Solver (Jo et al. 2008) was used to calculate the electrostatic potential map. All pictures were drawn using PyMOL v0.99rc6 (Schrödinger and DeLano 2020). Atomic coordinates of the crystallographic models have been deposited with the Protein Data Bank (accession codes 8AMG, 8AMI, 8AMJ, 8AMK, 8AML, 8AMM, 8AMN). X-ray diffraction images are deposited in MX-RDR database (Macromolecular Xtallograpy Raw Data Repsitory) (https://mxrdr.icm.edu.pl/) with appropriate DOI (Table S1).

### UV melting of oligonucleotides

UV thermal melting studies were performed according to a previous report (Rypniewski et al. 2016) on a Jasco V-700 spectrometer with a thermoprogrammer. The RNA oligomers were dissolved in a buffer containing 0.1 or 1 M sodium chloride, 20 mM sodium cacodylate pH 7.0 and appropriately 5 mM of BaCl_2_ or CdCl_2_ or SrCl_2_ or CaCl_2_ or CsCl. Each duplex was prepared in nine different concentrations in the range of 10^−5^ to 10^−6^ M. The concentrations of single-strand oligomers were calculated from a high temperature (>80°C) absorbance and single strand extinction coefficients approximated by the nearest-neighbor model. The UV absorption versus temperature was measured at 260 nm at a heating rate of 0.5 °C/min in the range 5–90 °C. The melting curves were analyzed, and the thermodynamic parameters were calculated using MeltWin 3.5.

### DSC melting of oligonucleotides

The RNA oligomers were dissolved in buffer I containing 0.1 or 1 M sodium chloride, 20 mM sodium cacodylate pH 7.0 and appropriately 5 mM of BaCl_2_, CdCl_2_, SrCl_2_, CaCl_2_ or CsCl. The samples were dialyzed overnight at 4 °C against buffer I in order to equilibrate RNA and the reference solutions. DSC experiments were carried out on a MicroCal PEAQ-DSC calorimeter (Malvern Instruments Ltd.) Each measurement was performed in five cycles of heating and cooling in the range of 2–110 °C and a scan rate of 1 °C/min. First, reference scans of buffer I were carried out to establish the instrument thermal history and to reach a near perfect baseline repeatability. Then, RNA samples were measured at concentrations ranging from 160–298 µM. The results were analyzed using dedicated software implemented by Malvern Instruments. The melting temperature Tm was calculated by applying a two-state model fitting.

## ACKNOWLEDGMENTS

This work was supported by the Institute of Bioorganic Chemistry, Polish Academy of Sciences.

